# Range expansion shifts clonal interference patterns in evolving populations

**DOI:** 10.1101/794867

**Authors:** Nikhil Krishnan, Diana Fusco, Jacob G. Scott

## Abstract

Increasingly, predicting and even controlling evolutionary processes is a sought after goal in fields ranging from agriculture, artificial intelligence, astrobiology, oncology, and infectious diseases. However, our ability to predict evolution and plan such interventions in real populations is limited in part by our understanding of how spatial structure modulates evolutionary dynamics. Among current clinical assays applied to predict drug response in infectious diseases, for instance, many do not explicitly consider spatial structure and its influence on phenotypic heterogeneity, despite it being an inextricable characteristic of real populations. As spatially structured populations are subject to increased interference of beneficial mutants compared to their well-mixed counter-parts, among other effects, this population heterogeneity and structure may non-trivially impact drug response. In spatially-structured populations, the extent of this mutant interference is density dependent and thus varies with relative position within a meta-population in a manner modulated by mutant frequency, selection strength, migration speed, and habitat length, among other factors. In this study, we examine beneficial mutant fixation dynamics along the front of an asexual population expanding its range. We observe that multiple distinct evolutionary regimes of beneficial mutant origin-fixation dynamics are maintained at characteristic length scales along the front of the population expansion. Using an agent-based simulation of range expansion with mutation and selection in one dimension, we measure these length scales across a range of population sizes, selection strengths, and mutation rates. Furthermore, using simple scaling arguments to adapt theory from well-mixed populations, we find that the length scale at the tip of the front within which ‘local’ mutant fixation occurs in a successive mode decreases with increasing mutation rate, as well as population size in a manner predicted by our derived analytic expression. Finally, we discuss the relevance of our findings to real cellular populations, arguing that this conserved region of successive mutant fixation dynamics at the wave tip can be exploited by emerging evolutionary control strategies.

## Introduction

Beneficial mutants and their substitution within a population encapsulate the very crux of natural selection, especially in its classic conceptualization. Using his now ubiquitous plots of genotypic frequency over time, H.J. Muller demonstrated how beneficial mutants can fix long before subsequent beneficial mutants arise, thus accumulating in a successive manner over time.^1,2^ Using a theoretical population genetics approach, J.H. Gillespie influentially termed these dynamics strong selection weak mutation (SSWM), after the relative balance between population size, mutation frequency, and selective advantage he derived for evolution under this model.^3–5^

Gillespie further demonstrated that evolution proceeding in this regime can be conveniently approximated using the robust mathematical construct of a Markov chain, in which evolution proceeds as an adaptive walk across a mutational landscape. Gillespie, and many others thereafter, have exploited the mathematical convenience of this regime to explore adaptation within molecular evolutionary data, devise algorithms to control the speed and direction of evolution, and examine the computational limits of evolution.^6–9^ More generally, this framework has proven a useful qualitative description for beneficial mutation and selection, as these ideas have been applied to experimental biology since Gillespie’s theoretical work.^10^. It is useful to be able recognize evolution under the SSWM regime given this Markov-chain representation of evolution, and the powerful analyses it allows (Box 1).

#### Box 1: Beneficial mutations in a well-mixed population

##### Beneficial mutant fixation and establishment times

For a large asexual population of fixed size *N*, with a large, finite number of beneficial mutations conferring fitness advantage *s,* acquired at rate *U_b_*, the mutant lineage population size, *n,* grows as 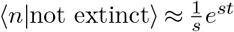 at long times, *t.* Thus we have the approximate average time for a mutant to establish, 〈*τ_est_*〉, and average time for a mutant to fix 〈*τ_fix_*〉,:

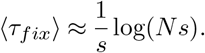

##### Dynamical regimes of beneficial mutations

When 〈*τ_est_*〉 ≫ 〈*T_fix_*〉, or *NU_b_* log(*N_s_*) ≪ 1, evolution is mutation limited, this is termed strong selection weak mutation (SSWM).

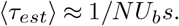

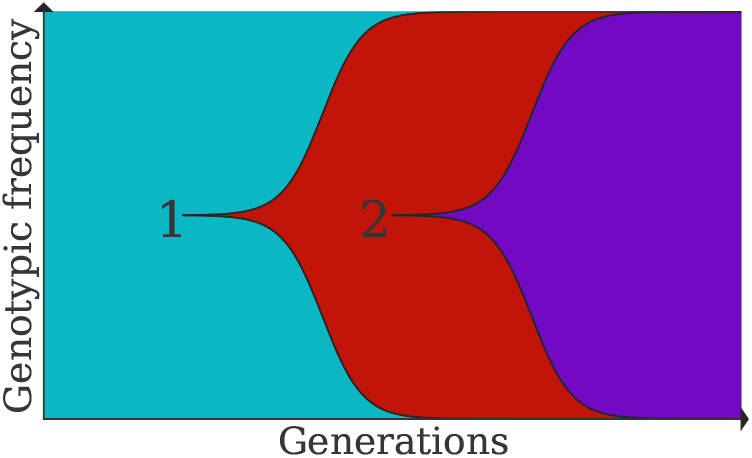

When 〈*T_fix_*〉 ≳ 〈*τ_est_*〉, *NU_b_* log(*N_s_*) ≳ 1, and clones are expected to interfere as they fix, this is termed strong selection strong mutation (SSSM). ^5^

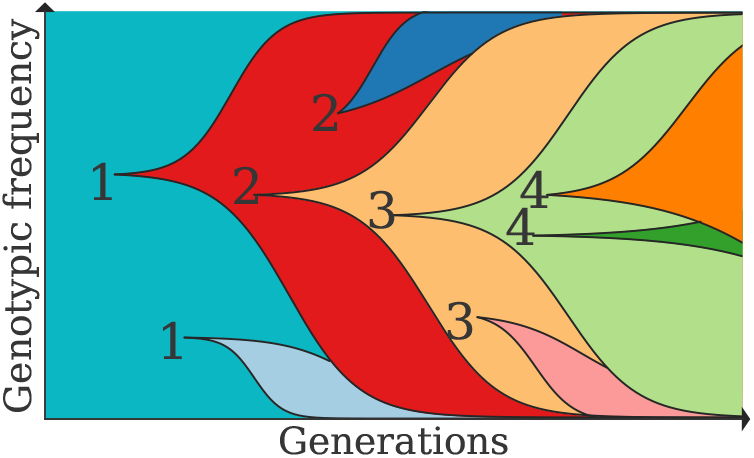

With even higher mutation frequency, the population is likely to have many coexisting mutations, with successful selection of any given mutant limiting the rate of adaptation. This regime can be characterized as weak selection strong mutation (WSSM).^10^

However, making such assumptions once spatial complexity is introduced presents a challenge. The spatiotemporal dynamics of mutants introduced in a migrating population can be surprising, complex, and resistant to analytical solutions. After spread into a new environment, the individuals at the very tip of the expanding population are preferentially represented in the resulting population’s genetic makeup, as observed across scales and domains of life.^11–13^ Relatedly, mutants at the front of a population expanding its range have an increased probability of survival and further proliferation at the wave front, a phenomenon known as gene surfing.^14^ Such results illustrate a fascinating positional dependence of evolutionary dynamics in spreading populations, an observation that has sparked many lines of investigation both experimental and theoretical.^15–18^ Great theoretical progress on population expansions has been made analyzing modelling frameworks based on stochastic reaction-diffusion systems.^19^ Recently this has included work on the properties of range expansions including range expansion with neutral mutations,^20^ range expansion in the strong noise limit,^21^ selection in the absence of mutations,^22^ long-range dispersals,^23^ and cooperation during range expansion.^24,25^

The interference of beneficial clones as they fix through a spatially extended population undergoing mutation and selection, however, represents an analytical challenge. The dynamics of clonal interference, even within well-mixed populations, are complex, and an active area of investigation in which recent advances rely heavily on simulation-based studies.^10,26^ In the case of spatially extended populations, Korolev et al. has shown for a one-dimensional stepping stone model with two alleles with unequal fitness benefits, and a non-zero mutation rate between them, a closed-form analytic solution to the spatiotemporal dynamics of the allele frequencies cannot be achieved without approximations that ultimately yield largely inaccurate results.^20^

Despite these limitations, Martens and Hallatschek have made progress in modeling the interference of beneficial mutants in a spatially structured population by considering low mutation rates and relatively large spatial scales.^27^ The authors show that during migrations across large habitats, waves of beneficial mutants arising on distinct genetic backgrounds are likely to interfere beyond a certain characteristic habitat length, determined by the frequency and strength of the mutations. Within this habitat length, one beneficial mutant wave fixes locally on average, while interfering with previous waves further from the edge of the mutant beyond this habitat length. We note that in this work Martens and Hallatschek refer to this as ‘clonal interference’, while this term, as originally coined by Gerrish and Lenski, refers to interference between two beneficial mutant arising in the same genetic background.^28^. In the language of Gerrish and Lenski, Desai and Fisher, and others, Martens and Hallatscheks’ work on the inference of slowly advancing waves of beneficial mutants with distinct genetic backgrounds during long-range migration is the ‘multiple mutation’ phenomenon, and hereafter we will refer to these separate phenomena using this convention.^5,29^

In this study we will build upon previous work showing dynamical patterns that emerge during the process of mutants arising and fixing within their local environment are positionally dependent with respect to origin along the population front of expanding populations. We employ agent-based simulations of a modified one-dimensional stepping stone model and analysis based off of the estimated time and length scales of mutant establishment and fixation. We demonstrate that within a co-moving frame, a stable sub-population at the tip of a spreading population front ‘locally’ fixes beneficial mutants analogous to a well-mixed population within the successive-mutant regime, while simultaneously, proximal regions of the wave front experience clonal interference. In addition, we discuss the biological relevance of our results as a whole.

### Model

Our model begins with a one-dimensional lattice configuration representing a linear habitat. Along the habitat, at each lattice site are colonization sites with a fixed capacity of *K* particles, called demes. Within each deme, these particles can either represent a vacancy or an individual, with some number of mutations. For each simulation step the following three steps are carried out:

1. A migration step in which two neighboring demes are chosen at random and a random particle from each deme is swapped with one another.
2. A duplication step, in which two particles are chosen from a randomly chosen deme, and the second particle is replaced by a duplicate of the first with some probability.
3. A mutation step, in which a random particle is granted a mutation with some probability.

These steps are schematized in Fig. 1. The probability of duplication during the duplication step is 1, except for the case of replacing an individual with a duplicate of a vacancy (i.e. death), which occurs with probability *1-r_m_*, where *m* is the number of mutations the particle has acquired. The transition probability of a non-vacancy particle acquiring one mutation during a time step is *U_b_* and 0 for all other possible mutational events (i.e. no backward or multiple mutations are allowed within a time step). The above model is akin to a modified stepping stone model used in previous literature and generates stochastic Fisher-Kolomogrov-Petrovsky-Piscounov (sFKPP) waves undergoing selection and mutation.^30^ The speed of the Fisher wave established from any individual is dependent upon the parameter *r_m_*, which can shown to be analogous to the growth rate of the corresponding sFKPP system when written in the continuous limit with appropriate assumptions (See SI, sec. 1.2.1).

**Figure 1.**
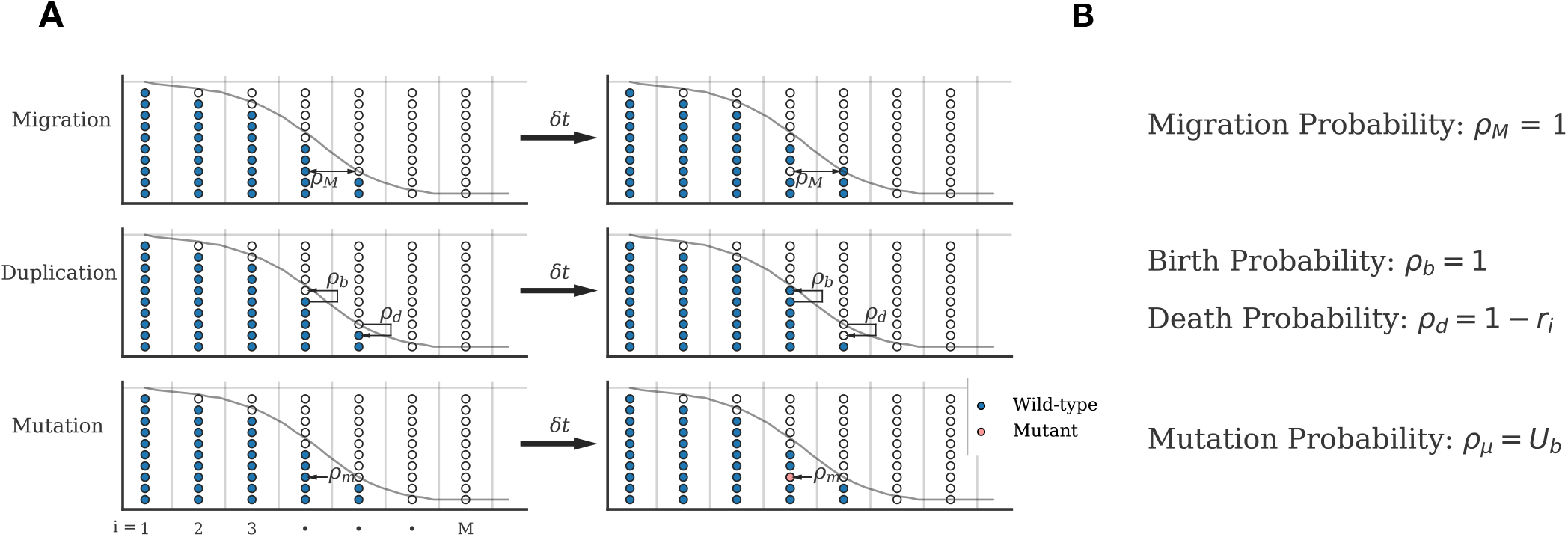
Simulation algorithm: one-dimensional stepping stone model with selection and mutation. A) The simulation field is comprised up to *M* demes distributed along a line, shown here along the horizontal axis. Each deme can each contain up to *K* particles, shown here as distributed along the y axis within each deme. Each particle is either a vacancy (white) or individual with *m* mutations, with, in this diagram, *m* = 0 (blue) or 1 (pink). The simulation is comprised of a Migration, Duplication, and Mutation step, illustrated from top to bottom respectively, which each occur once per time step *∂t*. Each of the 10 possible events is diagrammed with an arrow, illustrating the change in the system state per time step 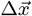, and its associated probability *p_E_*. The simulation is equivalent to a sFKPP wave in the continuous limit, arbitrarily re-scaled, and superimposed on the simulation field as a dotted curve. B) Each possible event and the corresponding probabilities with which they occur during a time step for a randomly chosen particle in two neighboring randomly chosen demes for Migration, two randomly chosen particles in a randomly chosen deme for Duplixation, and a randomly chosen particle in a random deme for mutation.

Simulations are initialized using a standing wave solution to the deterministic Fisher equation and we use an initial wild type wave of growth rate *r*_0_ = 0.1. As the cells spread throughout the linear habitat they undergo rare mutations that confer the organism a growth rate of *r_m_* = *r_0_α^m^,* where *α* quantifies the average strength of the fitness benefit conferred by the mutation, and *α* = 1 + *s* for *s* > 0. During the simulation, full demes furthest behind the wave tip are periodically dropped and empty demes are added past the wave tip. This choice improves computational efficiency, while allowing measurement of the evolutionary dynamics along the portion of the front, where mutant survival provability varies most significantly with position.

In this way, the simulation field represents the wave front, traveling at the average velocity of the spreading population. As mutants accumulate over time, their clonal sub-populations grow and establish into travelling waves with some probability, eventually causing the wild-type population to fall behind the mutant wave-front. Simulations stop at the extinction of a specified number of mutants within the the wave front, with the number of distinct clonal populations in each deme recorded. All simulations were performed in the python programming language with the Numba package to optimize computational speed.^31^

## Results

### Fate of a beneficial mutant

At short spatial and temporal scales and sufficiently large deme sizes, we estimate that the dynamics of mutant establishment are roughly as in a well-mixed population. To emphasize this analogy we define *s* ≡ *α* − 1. At sufficiently low mutational supply and strong selective mutant strength, a mutant establishes locally with probability ~ *s*, analogous to the ultimate fixation probability of a beneficial mutant in a well-mixed population of constant size (See Box 1).^5^ Accordingly, the reciprocal of this probability is the approximately average timescale of local establishment, *τ_est_* ~ 1/*s*. As shown by Lehe et al., the probability that a single mutant wave emerges from the wild-type wave after local establishment and goes on to outpace and take-over the wild-type wave (or surf on the wave front), in the absence of clonal interference, saturates at *r_0_* (1 + *s*) (conditioned on establishment) as the position at which the mutant is introduced goes further into the sparsely populated tip of the wave front (See Box 2).^14,22^ Before a mutant establishes, at a time *t* < *τ_est_*, it diffuses along the axis, and, if it fails to successfully establish, is left behind by the traveling population front. With sufficient mutational supply and strength, multiple mutants may arise along the wave front and interfere if their spatial scales of diffusion overlap or if one or more subclones establishes into a traveling wave and migrate to neighboring demes (See Fig. 2).

**Figure 2.**
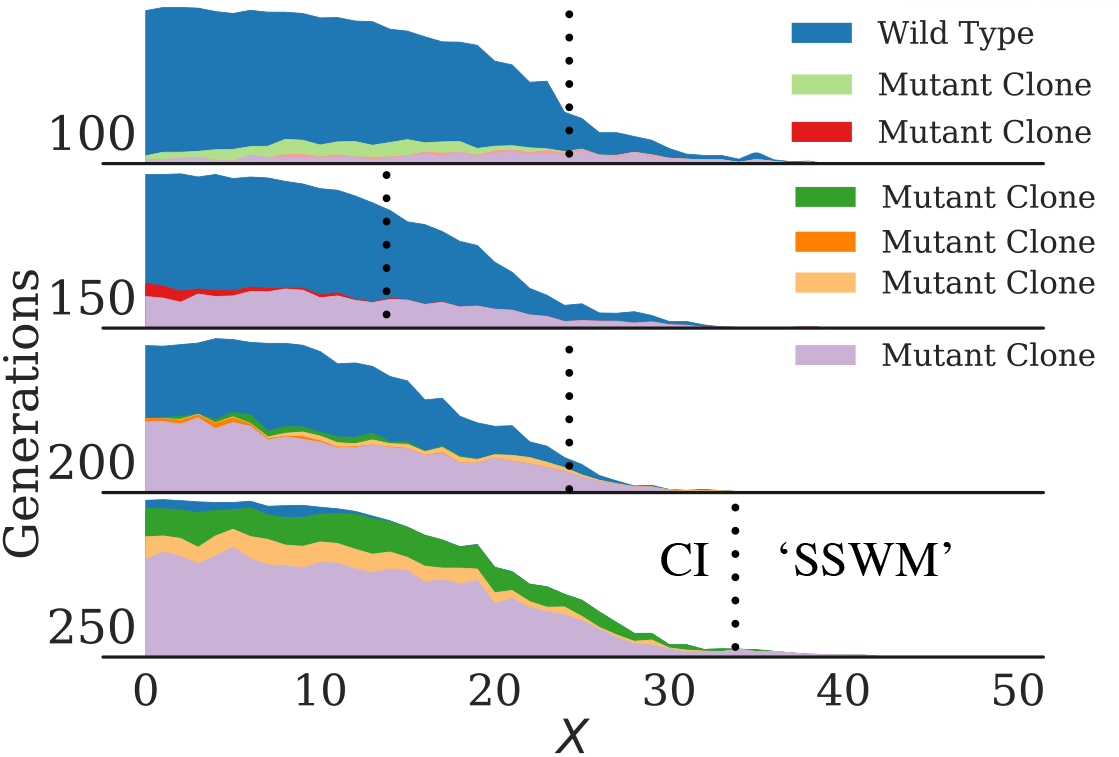
Mutant clones interfere with each other in various stages of diffusion, moving as an establishing traveling wave, and successfully surfing on the wave front. The wave profile in the co-moving frame of a population that can acquire beneficial mutations during range expansion is shown at a particular time point on each row, with increasing time from top to bottom. The y - axis represents the total population density while the x-axis represents the position *x* in the co-moving frame. Wild-type individuals are indicated in blue, and each other color a distinct mutant clonal population, each with the same selective advantage relative to its genetic background of origin. Mutant clonal populations are variously observed as traveling waves beginning to surf at the front of the population (violet), established mutants that have failed to surf at the wave front (red, generation 150), and spreading diffusively along the x-axis without apparently establishing as a traveling wave (salmon, generation 100). Dotted lines are placed at the position at which the population transitions from being polyclonal (labeled ‘CI’ for clonal interference) to monoclonal (labeled ‘SSWM’ for strong selection weak mutation) within local spatial scales. Simulations were performed with mutation rate *U_b_* = 10^−3^, selective advantage *s* = .1, and diffusion constant *K* = 500.

### Length scales of evolutionary regimes

In our initial simulations, the genetic diversity (number of distinct subclones) in each deme throughout the wave front was recorded at the time of wild-type extinction. From our simulation data, with sufficient mutation rate, deme size, and mutation strength, we observe a length scale extending from the very wave tip to some position within the bulk of the wave front in which the population remains genetically pure on average (Fig. 3). Following the scaling arguments of Martens and Hallateschek, we can quantify the time scales of the previously described scenario of mutant inference to illustrate the quantitative relationships between these parameters and this length scale, which we denote as *L_SSWM_*.^27^ A single beneficial mutant that has established into a traveling wave will fix in time *τ_fix_* by travelling across a length *L* such that

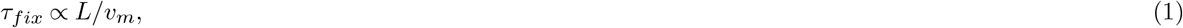

where *v_m_* is the relative velocity of the mutant wave. Within the length scale *L*, the average time for mutants to establish (*τ_est_*) locally is estimated by the following relation:

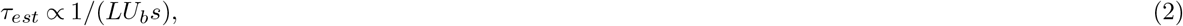

somewhat analogously to a well-mixed population (See Box 1). These times are comparable at a particular length scale at which mutants begin to interfere locally as they fix. We find *L_SSWM_* by setting *τ_fix_ ~ τ_est_*:

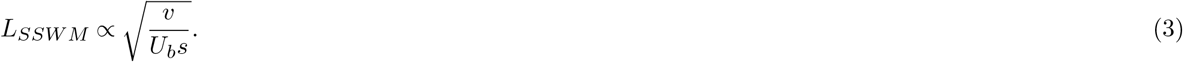

**Figure 3.**
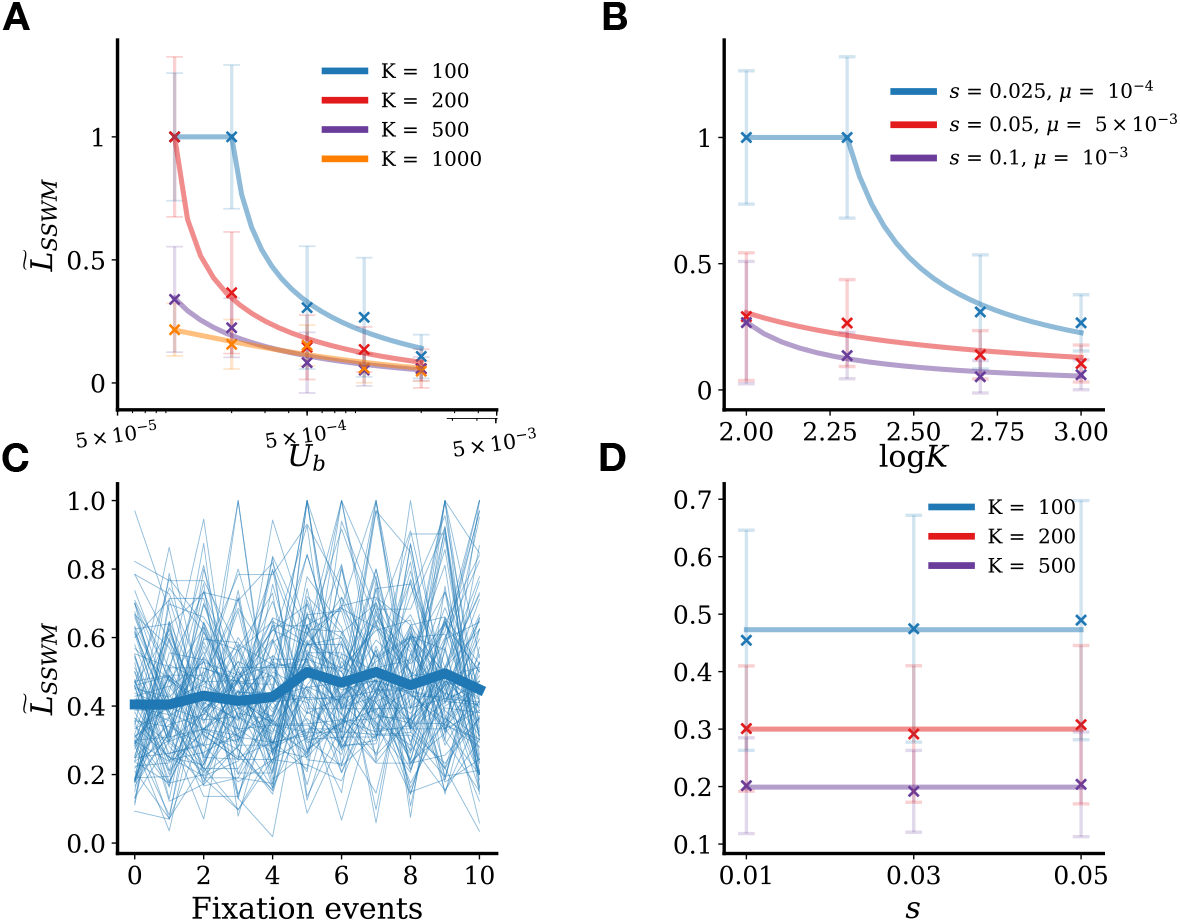
Proportion of wave front in ‘local’ successive mutation regime goes as 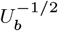 and log^−1^ *K*, and saturates at 1 with sufficiently weak mutational supply and strength. Over an ensemble of simulations (~ *40 forpanelsA, B;* ~ *150 forpanelsC,D*)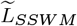 is measured as the distance in demes from the wave tip to the position at which no more than two clonal populations exist, divided by the total number of demes occupied by the wave front in the co-moving frame. **A)** Over a range of *U_b_* and *K* with *s* = 0.1, the median 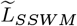 from simulation results and standard deviation (shown with X’s and error bars respectively are plotted showing approximately approximate agreement with an inverse square root fit (colored lines) well, as predicted by Eq. (3). 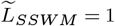 corresponds to successive mutant fixation throughout the wave front. **B)** 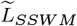 is again measured from simulations and plotted against log*K* at three particular values of *U_b_* and *s*, showing approximate agreement with an inverse fit, as predicted by Eq. (3). **C)** 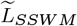 shown for each of 10 extinction events over 100 instantiations for s = 0.01, K = 100, *μ =* 10^−3^ show strong fluctuations, but remain roughly constant on average (thick blue line). **D)** Median 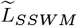 over 10 extinction events at *K* = 100,200,500 and *μ* = 10^−3^, 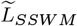 does not vary over *s =* 0.01,0.03,0.05. Simulation data is fit with line of slope 0 as predicted by Eq. (3) for relatively weak selection assuming a noisy Fisher wave velocity proportional to *s*.

Within this length scale, we expect local fixation of an established mutant to occur before another mutant establishes locally, on average, analogous to a well-mixed population within the SSWM regime.

Thus, from Eq. (3) we expect *L_SSWM_*, the length scale at the tip of the wave in which we expect mutants to appear successively to go as 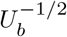 which agrees well with our stochastic simulations, see (Fig. 3A).

As expected, this behavior varies strongly with deme size. As the total length of the wave front, *L_front_*, defined as the furthest deme with greater than one individual, varies linearly with total population size, within the wave front we expect a linear relationship with log *K*. This can be demonstrated by noting that the initial wild-type population density at the tip of the front in the continuous limit *b_w_ (x)* (As in Box 2 Fig. L), can be approximated as an exponential decay function,

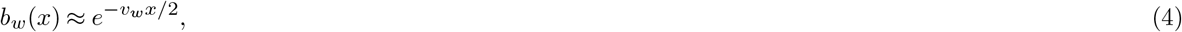

for *x >* 0 and *b_w_* (*x*) ≡ 1 everywhere else, with position *x* and velocity *v_w_*. As *x* for which *K* * *b_w_* (*x*) = 1 determinesthe length of the wave front, *L_front_,* solving Eq. (4), we obtain

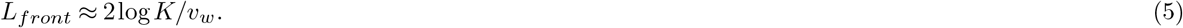

Equation (5) implies that *L_SSWM_* expressed as a proportion of the total front,*L_SSWM_*/*L_front_*, hereafter 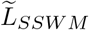 for simplicity, is inversely proportional to log*K*. This relationship is observed within our simulation data as well (Fig. 3B).

We additionally observe that with relatively weak mutational supply or mutation strength, this heterogeneity in evolutionary regime is not observed, and instead the entire wave front accumulates beneficial mutants successively. We estimate this to occur when the mutational supply relative to its selective advantage is sufficiently small: 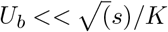 When this constraint holds, it is expected that 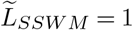 on average throughout the wavefront (See Supplementary Information, section 2.3).

##### Box 2: Mutant surfing on the wave of an expanding population

###### Beneficial mutant surfing probability

Mutant surfing probability with respect to position, *u(x)* is well approximated as a stochastic branching process with a phenomonoligical correction (*u*^2^ term):

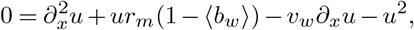

The balance between increased surfing probability at the wave tip, and larger mutational supply toward the wave bulk can result in the surfing mutant supply being maximized somewhere in between (See SI sec. 1)

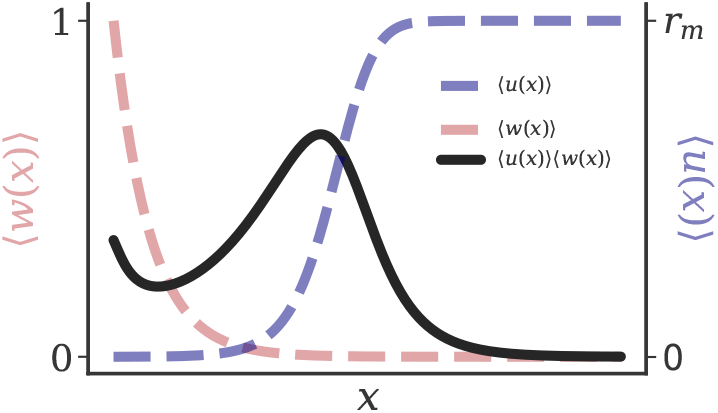

###### Beneficial mutant substitution rate

The supply of successful surfing mutants can approximately be quantified as:

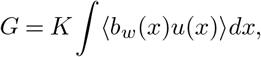

and the substitution rate for the population front is given by *GU_b_*. In a well-mixed population the analogous expression to *G* is represented as *K_s_*, for total population size *K* and selective advantage *s*. Compared to a well-mixed population, where the substitution rate increases weakly with *s*, especially at low values, demonstrates that selection is rendered inefficient by range expansion.^22^

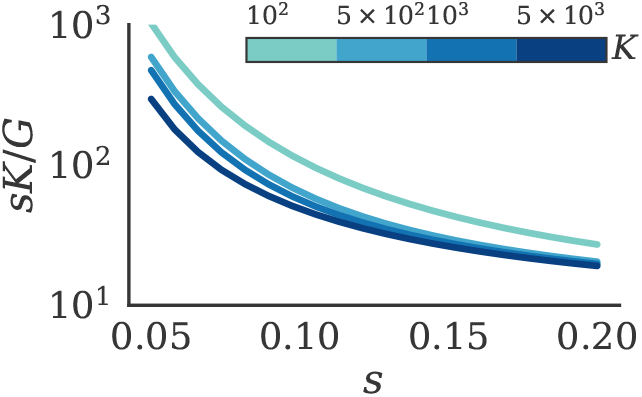

To further examine the relationship between *s* and *L_SSWM_*/*L*, we simulated evolution in our system through multiple extinction events and observe that if the evolutionary regime switches at some position in the wave tip, this regime switching is sustained and roughly invariant over multiple extinction events and across evolution scales(Fig. 3C). As each mutation event on average occurs at a particular time in the ‘cycle’ of mutant establishment and fixation (which parameter specific), 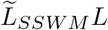 is seen to vary stochastically over time, though predictable relationship between 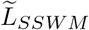 with *s* is observed over an average of many extinction events. Our simulations of multiple extinction events were limited by the maximum fitness at the end of simulations, which we ensured obeyed our analytic assumptions, i.e. *r_m_* ≪ 1 by performing these simulations at low values of *s* ∈ 0.01,0.03,0.05. Multiple fixation events at higher values of *s* or *K* would allow for mutants that violate the assumption of strong migration 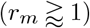, under which our analysis holds.

The relationship between 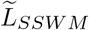 and the selective advantage at each extinction event in our simulation is more nuanced as *s* determines mutant establishment probability, velocity, and the shape of the front profile. While the velocity *v* for a classic Fisher wave 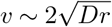 with growth-rate *r*, there are two important deviations from this classic behavior.

First, as waves interfere with each other, as is the case within the parameter regime we simulated, Martens and Hallatschek demonstrated that their velocity slow with increasing interference, though this velocity does saturate at a critical value. Secondly with decreased *s* relative to the strength of the noise, the velocity deviates from the deterministic velocity 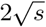, and becomes roughly proportional to the product of *s* and wave density, as opposed to 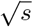 in the deterministic limit ^21^. These two phenomenon underscore the complexity in the relation between *L_SSWM_* and *v* and *s*. At strong noise relative to *s* as in the parameter regime simulated, for example, we expect *v ∝ s* and thusly 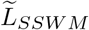 to be invariant with *s*, from equation (3). Simulation results from *s* = 0.01,0.03, .05, *K* = 100,200,500 and *μ* = 10^-^3 show 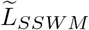 to be roughly invariant with *s* in this regime. (Fig 3D)

## Discussion

An enduring goal of population genetics is to understand the temporal dynamics of genotypic frequency and the selective forces that (may) shape them. When mathematical analyses or simulation models are used for these means, it is important, then, to consider the extent to which our model does and does not reflect known biology to be able to evaluate the validity of the results it provides.

To this end, we first consider the range of experimentally observed beneficial mutation rates. Beneficial mutation rates have historically been posited to be very low, and early experiments supported this with estimates on the order of 10^-9^ − 10^-8^ per genome per generation.^28,32^. However, subsequent experiments that have sought to eliminate the bias introduced by non-optimized growth conditions, and unaccounted for clonal interference, variously suggest beneficial mutation rates may be on the order of 10^-^4. Accurately measuring and even identifying mutations as beneficial in an empirical context is still an active area of investigation.^28,32^ Our model examines parameters an order of magnitude above below this upper limit.

We secondly consider the strength of beneficial mutations observed previously. Recent studies have been in closer agreement with measurements often from .01 − 0.05. In our study, we examine this range of selective advantages within and up to an order of magnitude above the empirical values.

Thirdly, we address deme size in our model and its relation to real populations. We note that in a one dimensional stepping stone model, deme size is effectively a linear density term, however its relation to real population is not straight-forward. In this study, as in others we explore a relatively wide range of deme size to capture effects specific to any given regime. However we can attempt to gain a rough idea of the relationship of deme size to population density of real populations by stating deme size in terms of maximum individuals per diffusion length, or average distance diffused within a generation. Given *D* = 1 and *r_0_* = 0.1, in our model 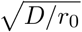 describes a diffusion length, and 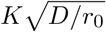 the maximum number of individuals within a diffusion length For values of *K* ranging from 100 to 1000 as examined in our simulations this corresponds to ~ 300 − 3000 individuals per diffusion length.

To compare this estimate to real cellular populations, we can take the example of bacterial colonies, which are observed to exhibit diffusive motility within a large range of densities.^33^ To get a sense for this range we refer to Butler et al. and Perfeito et al. who report report a migrating cell surface density of ~ 3 × 10^6^*cells/mm^2^* on agar plates. We further use and average generation time (i.e. experimental *r_0_* of .7 *hr^-1^*) and a range of cell diffusivities from 0.7 − 5.7 mm/hr, observed with varying agar concentration for *E. Coli.^34^* We can further suppose a minimum possible surface density of close packed *E. Coli* without deformation as 1 cell per *μm^2^*. Converting the experimental surface densities to a linear density *ρ*, we find individual per diffusion length as 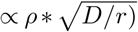.

This yields a range of 1000 to 5000 cells per diffusion length and overlaps significantly with the estimated linear density examined in our model, suggesting the model parameters examined and the corresponding behavior observed are biologically plausible. Furthermore, at higher densities than those examined in our model, shifting clonal interference patterns within a wavefront can be observed at even lower mutation rates (See SI 1.2.3).

The aforementioned discussion suggests a biological relevance for interference with relatively high beneficial mutation rate and short spacial scales as examined in our model. We have shown that in this regime within a relatively wide range of biologically relevant parameters, a region at the tip of the wave front fixes mutations ‘locally’ in a successive manner, analogous to the SSWM regime in well-mixed populations. Such a property is potentially useful for applying analyses such as a mutational landscape model, or considering theoretical work built on models assuming SSWM regime for the crucial tip of a wave expanding its population. In a variety of contexts, as the population ‘pioneers’, the individuals at the tip of the population and their dynamics can be considered exclusively, as they disproportionately influence the genetic diversity of the population following migration. In addition, our results demonstrate how the effective evolutionary parameters of a population shift through range expansion. The implications of the results for inferring evolutionary dynamics, for antibiotic resistance assays in clinical contexts, for instance, is that the dynamics observed at a specific time and place in the population maybe transient and highly unrepresentative of the entire population. While the spatiotemporal dynamics of a population expanding its range and acquiring mutations subject to selection can be highly complex, evading exact analytic treatment, we believe the our work demonstrates that analyses from well-mixed populations can be applied to the tip of the wave as long as it is maintained within the SSWM regime. To this end, we have estimated, in terms of ‘real parameters’ when this phenomenon is observed and the dependence of this scale length of local successive mutation fixation on relevant population genetic parameters. In summary, we shed light on the phenomenon of shifting clonal interference patterns during range expansion with relevance and application across evolutionary and scientific contexts.

## Acknowledgements

The authors would like to thank Dr. Diana Fusco for her thoughtful feedback and discussion. JGS would like to thank the NIH Loan Repayment Program for their generous support and the Paul Calabresi Career Development Award for Clinical Oncology (NIH K12CA076917). He would also like to thank the NIH for their support through NIH399R37CA244613 and the American Cancer Society for the Research Scholar Grant (Award number: 132691- RSG-20-401096-01-CSM).

## Author contributions statement

NK performed the mathematical analysis, wrote the code, performed the simulations, analyzed the data and wrote the manuscript. JGS analyzed the data and wrote the manuscript.

## Code availability

The code to perform the numerical simulations is available via github at https://github.com/nkrishnan94/Range-Expansion-Eco-Evo-Regime. https://www.ncbi.nlm.nih.gov/pmc/articles/PMC2975585/

## 1 Supplementary Information

### 1.1 Analysis of beneficial mutant surfing

Here, we will review the analysis of the beneficial mutant surfing probability by Lehe et al., shown in Box 2 of the main text. In their work Lehe and colleagues demonstrate that the beneficial mutant surfing probability with respect to position of origin is well approximated as a branching process. Such a branching process yields the following differential equation with a non-linear correction term.

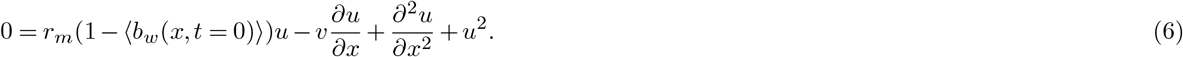

The analytic solution *u(x)* can be expressed in terms of a length scale *γ*, which governs how far in the wave tip the surfing probability begins to rise significantly.

In Box 2, to demonstrate how the mutational supply for surfing mutants is non-monotonic with respect to position we follow Lehe et al: the initial wild type wave *b_w_*(*x*) can be approximated as 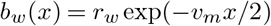 The mutant surfing probability can be approximated as *u(x)* = *r_m_*exp(*v_m_*(*x* − *γ*)/2), where 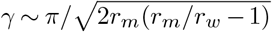. Using this approximation, one can find 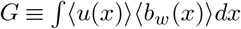, for sufficiently strong selection, which allows for an approximation of the substitution rate given the initial wild type wave density and survival probability. Using the above approximate expressions for *u(x),w(x)* Lehe et. al. further find an analytic expression for *G* in the limit of large *K* and and small *s*, though this expression and analysis was not used for the demonstration in Box 2

### 1.2 Mathematical expression of simulation algorithm

#### 1.2.1 Master equation

We consider demes arranged along a single dimension denoted by *i* =1,2,3… *M*. The vector 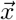 describes the number of individuals 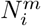, with *m* ≥ 0 mutations at each deme i from 1 to 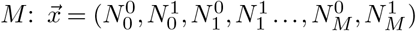 Over the course of a time step, the events that occur with the simulation are denoted by the index *E*, its respective probability as a function of the state vector *pE*(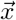, and the change in the state vector 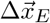. We can then describe change in probabilities for the entire system, 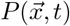 over a time step within our algorithm with the following general master equation:

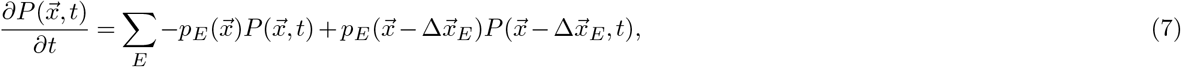

where *p_E_* and 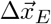 for all E are as follows:

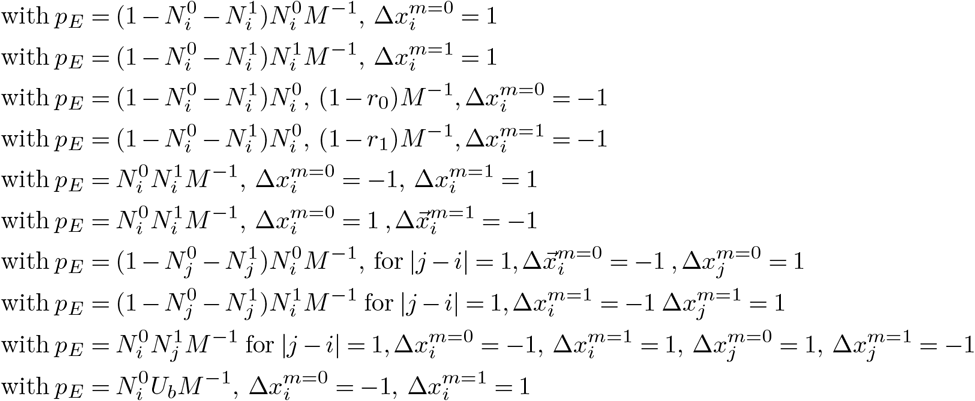

#### 1.2.2 Continuous limit of model dynamics as Stochastic Fisher-KPP equation

To discuss the examine the bounds of our model validity one apply the diffusion approximation to the master equation (7) obtaining a Fokker-Planck equation and subsequently the corresponding Langevein equations, corresponding to a stochastic Fisher-KPP equation with mutation (See Refs. ^19,20,22,35^):

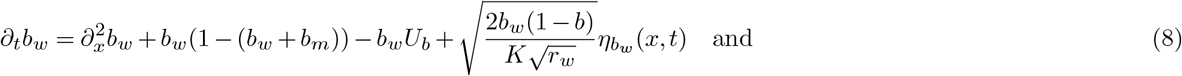

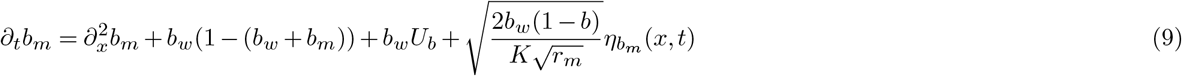

where *b_w_ (x,t)* and *b_m_ (x,t)* are the wild type and single mutation population densities through time and space (non-dimensionalized) respectively. For simplicity we omit the populations carrying multiple mutation and only consider the wild-type and single mutant populations. *r_w_* and *r_m_* describes the average growth rates per unit time and *Ub* is the rate at which wild type individuals acquire beneficial mutations per generation

Equations (8) and (9) are valid assuming 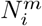 is sufficiently smooth across i, *r_m_,r_w_* ≪ 1, and *η_b_w__ (t),η_b_w__* (*t*) are Gaussian white noises uncorrelated with itself at each *x* and *t*.

#### 1.2.3 Bounds on model validity

From the above description we can analyze the boundaries in terms of model parameters at which our results our expected to be valid following the analysis of Martens and Hallatschek.^27^ Initially mutations are expected to occur at an average time *τ_mut_ ~ 1/UbK*.

In analogy to the heuristic arguments for a well mixed population (Box 1), we expect a mutant to establish on time scales ~ 1/*s*. In that time the mutant population will diffuse over a length 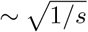, given a migration rate of 1 by construction of our model. In regions where mutants are most likely to interfere during this phase the local population within this length scale is roughly 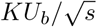 Approximating the time between mutant establishment on this scale as 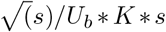, interference is unlikely when the time over which mutant establishment occurs ~ 1/*s* is much less than time between mutant arrivals. Stated mathematically:

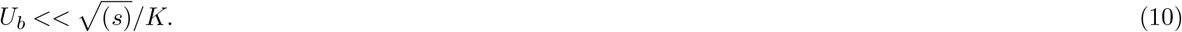

In the context of our model, *s* = *α* − 1. We can develop a sense of the validity of this predicted constraint by considering *U_b_* below 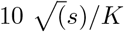 to be sufficiently in accordance with Eq. (9). Fig. 3D in the main text shows this relationship plotted in parameter space with the simulations performed. The simulations that do not display this heterogeneity in clonal interference pattern fall roughly below the plotted surface 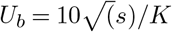. While a factor of 10 is an arguably arbitrary quantification of this constraint, these results suggest this constraint to be valid. Of note, by modeling mutations far below this upper bound, Martens and Hallatschek model the dynamics of this system with an algorithm in which each agent represents monoclonal subpopulation traveling as an established wave. ^27^.

